# Active reconfiguration of cytoplasmic lipid droplets governs migration of nutrient-limited phytoplankton

**DOI:** 10.1101/2021.10.17.463831

**Authors:** Anupam Sengupta, Jayabrata Dhar, Francesco Danza, Arkajyoti Ghoshal, Sarah Müller, Narges Kakavand

## Abstract

As open oceans continue to warm, modified currents and enhanced stratification exacerbate nitrogen and phosphorus limitation, constraining primary production. The ability to migrate vertically bestows motile phytoplankton a crucial–albeit energetically expensive–advantage toward vertically redistributing for optimal growth, uptake and resource storage in nutrient-limited water columns. However, this traditional view discounts the possibility that the phytoplankton migration strategy may be actively selected by the storage dynamics when nutrients turn limiting. Here we report that storage and migration in phytoplankton are coupled traits, whereby motile species harness energy storing lipid droplets (LDs) to biomechanically regulate migration in nutrient limited settings. LDs grow and translocate–directionally–within the cytoplasm to accumulate below the cell nucleus, tuning the speed, trajectory and stability of swimming cells. Nutrient reincorporation reverses the LD translocation, restoring the homeostatic migratory traits measured in population-scale millifluidic experiments. Combining intracellular LD tracking and quantitative morphological analysis of red-tide forming alga, *Heterosigma akashiwo*, along with a model of cell mechanics, we discover that the size and spatial localization of growing LDs govern the ballisticity and orientational stability of migration. The strain-specific shifts in migration which we identify here are amenable to a selective emergence of mixotrophy in nutrient-limited phytoplankton. We rationalize these distinct behavioral acclimatization in an ecological context, relying on concomitant tracking of the photophysiology and reactive oxygen species (ROS) levels, and propose a dissipative energy budget for motile phytoplankton alleviating nutrient limitation. The emergent resource acquisition strategies, enabled by distinct strain-specific migratory acclimatizing mechanisms, highlight the active role of the reconfigurable cytoplasmic LDs in guiding vertical movement. By uncovering the mechanistic coupling between dynamics of intracellular changes to physiologically-governed migration strategies, this work offers a tractable framework to delineate diverse strategies which phytoplankton may harness to maximize fitness and resource pool in nutrient-limited open oceans of the future.

**One sentence summary:** Phytoplankton harness reconfigurable lipid droplets to biomechanically tune migratory strategies in dynamic nutrient landscapes.

## Introduction

Oceans, today, are undergoing a major makeover due to the warming temperatures and perturbation of the currents, impacting the concentrations and availability of key nutrients across scales (*1–3*). High temperatures promote stratification of the ocean column, exacerbating nutrient limitation in tropical and mid-latitude surface waters (*4*, *5*). Together with the increased concentration of atmospheric carbon dioxide that trigger indirect changes of the surface water chemistry (*1*), the limitation of nitrate (NO_3_^−^), phosphate (PO_4_^3−^), and iron has impacted primary productivity in vast swathes of open oceans (*6*). The restructuring of the mixed layer depth exert critical control on the vertical distribution of phytoplankton (*7*, *8*), and associated community scale structure and interactions (*9–11*). Nutrient limitation, an inescapable fate of future oceans, is projected to continue amplifying uninterrupted, mediated by nutrient trapping (*5*), stronger stratification (*12*), and co-limitation of one nutrient by another limiting nutrient (*13*, *14*), with far-reaching ramifications on all major marine ecosystems (*2*, *4*, *5*).

Phytoplankton, key photosynthetic microorganisms occupying the base of nearly all aquatic food webs, acclimatize in the emerging nutrient landscapes by repositioning along the vertical column (*11*, *15–18*), aided by a range of coping mechanisms, both behavioral and physiological. Under diffusion-limited settings, the termporal nature (stable vs time-varying limitation), specificity of nutrient affinity, and species-specific allometries of uptake, maximal growth rates and nutrient storage, vary along the ocean column (*2*, *19–22*). Altered cell size (*23*), enhaced nutrient affinity (*24*), higher stoichiometric plasticity (*25*), and adjusted energy storage capacity (*26–28*), are among critical trait shifts which allow phytoplankton to redistribute along the ocean depths to maximize fitness. High nutrient affinity alongside low storage capacity offer competitive advantage to small-sized cells exposed to steady or frequently-pulsed low nutrient concentrations, while larger sizes are beneficial in oligotrophic waters if the cellular nutrient quota remains steady, as evidenced in vacuolated phytoplankton (*29*). Complementarily, maintaining a steady size can be advantageous for cells which can maximize nutrient affinity by down-regulating metabolically expensive processes, or by replacing key elements by widely-available surrogate elements in costly nutrient-specific cellular processes (*14*, *30*, *31*).

The ability to migrate, either by swimming or by buoyancy regulation, allows phytoplankton a critical access to nutrient-rich patches, as well as increase the flux of nutrient molecules (relative to pure diffusion), thus enhancing the competitive advantage to migrating species under low nutrient settings (*21*, *32*, *33*). Leveraging migration along the vertical ocean column, phytoplankton may explore a viable acclimatization alternative to the non-migratory trait shifts, provided the benefits outweigh the energetic costs. Whether a population migrates to optimize its resource searching strategy under nutrient constraints is contingent to the stored energy levels, along with the individual uptake capabilities should there be chanced encounters with ephemeral nutrient patches (*34*, *35*).

Energy-rich lipid and starch bodies allow phytoplankton to maintain fitness by gradual utilization of the stored resources (*26–28*). Over prolonged durations of nutrient limitation, heavier, short-lived starch bodies transform into lighter cytoplasmic lipid droplets (LDs). Though LD synthesis and maintenance come at the cost of immediate growth and division, LDs facilitate key metabolic processes during nutrient limitation, including crucially, amelioration of physiological stress (*36–38*). Larger storage capacity is potentially beneficial when nutrients are scarce (*39*, *40*), however migration can offset competitive advantages therefrom by enhancing encounters and expanding the available resource pool (*35*). NO_3_^−^ depletion has been associated with the initiation of diel vertical migration during red tides (*41*, *42*), enabling *Heterosigma akashiwo* access to the dissolved nutrients below the thermocline (*43*), yet contrasting observations with other species have indicated inverse correlations of lipid volume and phytoplankton motility (*44–46*). The jury is thus still out on the mechanistic links between cell-level LD dynamics and population-scale migratory shifts. The emerging conundrum that nutrient-limited phytoplankton face–in risking fitness due to limited growth and division, while enhancing migration-driven nutrient encounters–relies on the dynamics of storage and utilization of the energy-rich bodies. Yet how phytoplankton arrive at an optimal storage-and-migration strategy, a critical coupled trait, remains unknown.

Here we report that motile phytoplankton under nutrient limitation harness energy-rich cytoplasmic LDs to govern their migratory behaviour, expanding their resource pool and acquisition strategies. Using strains of *Heterosigma akashiwo*–a model gravitactic raphidophyte known for their toxic blooms (*33*, *47*) and rapid adaptability (*48*, *49*)–we show that cytoplasmic LDs nucleate as scattered droplets when concentrations of NO_3_^−^ and PO_4_^3−^ in the cell culture drop below a threshold value. Growth stages in our cell cultures, a proxy of ecologically relevant nutrient levels, trigger augmentation of LD size, followed by droplet merging and reduction in their count. Simultaneously, the expanding LDs translocate–directionally– through the cytosol to accumulate below the cell nucleus, triggering progressive modification of the ballisticity and stability of the swimming cells. We develop a data-based model of cell mechanics that establishes a mechanistic link between the LD dynamics and phytoplankton migration in nutrient limitation, elucidating the role of LD size and intracellular position in precisely governing the migratory behaviour. Nutrient reincorporation reverses the LD translocation restoring the swimming properties, highlighting an active control of phytoplankton migration by reconfiguration of the LDs. Together with observations of cellular stress levels, photophysiological performance, and selective switching to mixotrophy, our results point to an LD-governed migratory adaptation of phytoplankton under nutrient limitation, in a manner that accounts for the strain-specific differences we observe in the modified swimming strategies. We argue, using a dissipative energy budget, that storage and migration diversify as coupled traits in motile phytoplankton under nutrient limitation, allowing them to harness LDs biomechanically to regulate migration, thereby expanding the resource pool and diversifying the nutrient aquisition strategy. The amenable trade-offs between cellular behaviour and physiology we report here may ultimately equip species to maximize fitness in chaging nutrient landscapes of future oceans.

## Results

We track the growth and translocation of cytoplasmic LDs in two different strains of *H. akashiwo*, CCMP3107 and CCMP452 (hereafter strains S-1 and S-2 respectively) using cell-level quantitative imaging, alongside spectrophotometry to measure the concentrations of NO_3_^−^ and PO_4_ ^3−^ in the liquid cell cultures (Figure 1A, Methods). PO_4_^3−^ depletion occured faster, within ~125 h of inoculation, corresponding to the exponential growth stage, while NO_3_^−^ levels deplete below detectable values over a much longer period (*t* ~225 h, coinciding with the transition to stationary growth stage). The yellow-orange regions, initially scattered across the cytosol, represent intracellular LDs in S-1 (false-colour epifluorescence micrographs, Figure 1A, Methods), undergo significant growth as the NO_3_^−^ availability drops to *C* ~0 μM, 225 h after cells were freshly inoculated with *C_0_* = 600 μM of NO_3_^−^ (~165 pmole/cell). LDs do not nucleate till *t ~*200 h (Figure 1B), suggesting it is the sustained limitation of NO_3_^−^, and not PO_4_ ^3−^, that drives the growth of the LDs (*50*). Over the course of population growth, the cell area increases, reaching a maximum at the onset of the stationary growth stage (~180 μm^2^ at *t* = 396 h), coinciding with ~0 μM availability of NO_3_^−^. Thereafter the cell size reduces as the population enters the late stationary stage (*t* > 700 h). Correspondingly, the size of the LDs relative to the cell size, *I*_*LD*_ (*I*_*LD*_ = total lipid area/cell area), increases as the NO_3_^−^ availability drops, showing a significant enhancement when the population enters the stationary growth stage (ANOVA: P < 0.001; asterisk presents statistical difference in the lipid volume, *V*_*LD*_, inset Figure 1B). During the transition into the stationary growth stage (228 h < *t* < 300 h), *V*_*LD*_ increases 270%, at a rate of 0.12 μm^3^/h, thereafter maintaining a stable volume of ~20 mu^3^ into the late stationary stage (*I*_*LD*_ ~0.07 during the same period). Beyond *t* > 800 h, though *V*_*LD*_ remains stable (inset Figure 1B), *I*_*LD*_ increases further due to the reduction in the cell size, underscoring the simultaneity of distinct trait shifts–accumulation of energy-rich LDs and enhancement of the surface-to-volume ratio–in nutrient depleted populations (*23*).

**Figure 1.**
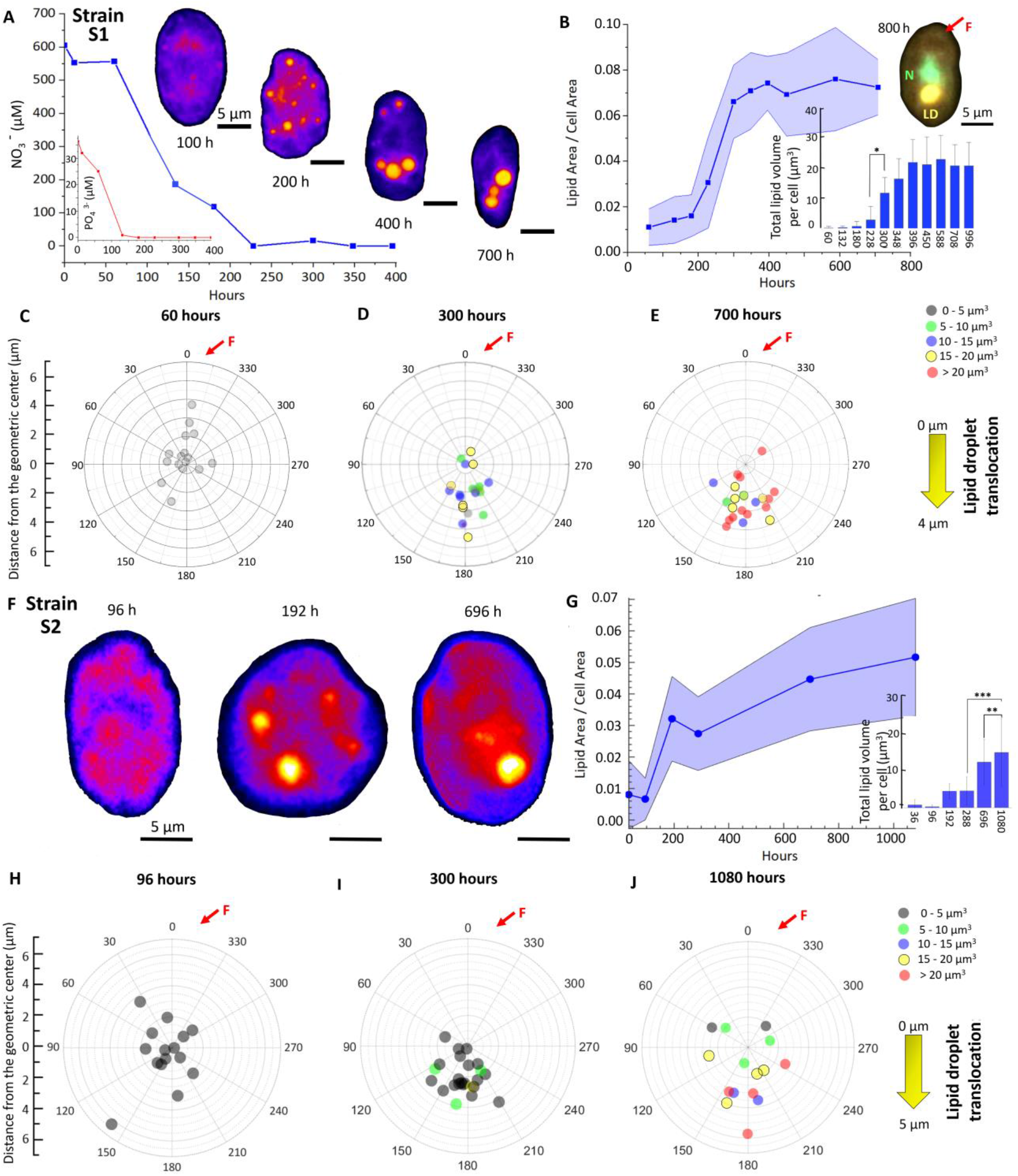
Nutrient depletion triggers cytoplasmic lipogenesis and lipid droplet translocation in phytoplankton. **(A)** Lipid droplet (LD, bright orange false-colour epifluorescence signal) nucleation, growth and translocation over time, triggered due to diminishing concentrations of NO_3_^−^ and PO_4_3^−^ (inset) in S-1 culture. Small LDs, scattered throughout the cytosol, are first visualized at 200 h after cell inoculation, as the population enters the late exponential growth stage. With the nutrient concentrations depleting further, reaching values below the detection limit of the spectrophotometer, the cell culture reaches its carrying capacity. In this nutrient depleted setting, the number of LDs per cell reduce, however the LDs grow significantly in size, and translocate within the cytosol, ultimately localizing below the cell nucleus, as visualized in panel B (top inset). **(B)** The normalized lipid size, *I*_*LD*_ (*I*_*LD*_ = total lipid area/cell area), increases steeply as the population enters early stationary growth stage (200 h - 350 h), and thereafter remains stable (*I*_*LD*_ ≈ 0.07) through the late stationary stage, before going up again after *t* = 800 h during which the cell reduces in size while the LD maintains a constant dimension. Squares and shaded regions denote mean ± s.d. of analyzed cells (*t*_60_ = 20, *t*_132_ = 20, *t*_180_ = 21, *t*_228_ = 21, *t*_300_ =25, *t*_348_ = 30, *t*_396_ = 21, *t*_450_ = 23, *t*_558_ = 24, *t*_708_ = 22). Top inset: A multichannel fluorescent micrograph shows localization of the LD (yellow-orange hue) below the nucleus (*N*, light green hue) in a nutrient-limited phytoplankton (*t* = 800 h). The relative location of the propelling flagellum (*F*) is indicated by the red arrow head. The bottom inset presents the *V*_*LD*_, the total lipid volume (sum of single LD in each cell) per cell, over time as a bar plot of the mean ± s.d.; ANOVA: P < 0.001; asterisk indicates significant difference in *V*_*LD*_ between *t* = 228 h and *t* = 300 h. *V*_*LD*_ remains stable at ~20 μm3 through all *t* > 400 h time points measured here. **(C-E)** Spatio-temporal coordinates of the effective LD in individual S-1, relative to the geometric center (or the center of buoyancy, *C*_*B*_) located at the center of the angular plot. *C*_*B*_ coincides with the cell nucleus and thus, with the cell’s center of gravity (*C*_*G*_). Tracking the effective LDs over time captures the intracellular translocation with depleting nutrients, i.e., with increasing age of the phytoplankton culture, over *t* = 60 h **(C)**, 300 h **(D)**, 700 h **(E)** respectively. The effective LD size, angular position, and the radial offset distance from *C*_*B*_ are obtained in individual cells, with their long axis indicated by the 0°-180° line (fore-aft direction). The fore and aft directions are distinguished by the position of the propelling flagellum located on the fore-end of the cell, slightly offset from 0° (shown as red arrow head, *F*). As indicted by the color-coded circles, the effective LD progressively moves toward larger size ranges, simultaneously translocating in the aft direction (away from the flagellum) with increasing radial offset from *C*_*B*_. **(F)** LD biogenesis and localization in S-2, shown here for *t* = 96 h, 192 h, 696 h. **(G)** The S-2 *I*_*LD*_ shows an increasing trend through the entire growth stage studied here (till *t* = 1080 h), reaching a maximum value of *I*_*LD*_ = 0.052 ± 0.018. Inset: Time series of *V*_*LD*_ for S-2 shows a significant increase from *t* = 288 h, as the cell culture reaches carrying capacity. t-test between 696 h and 1080 and one-way ANOVA between 288 h, 696 h and 1080 h, p < 0.001; asterisks indicate statistically significant difference. **(H-J)** Spatio-temporal coordinates of the effective LD in individual S-2 at *t* = 96 hours **(H)**, 288 hours **(I)**, 696 hours **(J)** reveal the growing size of effective LDs and concomitant aft-ward translocation of LDs over prolonged nutrient depleted conditions (see caption for panels C-E for quantification steps). Throughout the figure, the time axis and values denote the duration elapsed since the start of a fresh inoculum (nutrient replete condition), signifying the nutrient corresponding availability.

Phytoplankton regulate LD biosynthesis dynamically as the nutrient availability changes. Populations incoluated with lower initial nutrient concentration produced LD more rapidly (0.09 μm^3^/h in 1% C_0_ vs 0.076 μm^3^/h in control condition), though their LD carrying capacity, *i.e.*, the maximum volume of LD generated per cell, was lower compared to the control case (with 1% *C_0_*, *V*_*LD*_ ~10 μm^3^ < 22 μm^3^ for the control population). The reduced initial nutrient concentration, additionally, lowered the cell count over the entire growth stage, while enhancing the rate at which the cell size increases during the early exponential stage by nearly 45%. Logistic fitting of the LD volume per cell reveals that lipid accumulation ceases during the stationary growth stage, indicating phytoplankton do not consume LDs at the corresponding timescales. Taken together, our results demonstrate that, phytoplankton can adapt cell growth to accommodate the lipid yield and production rate dynamically, in face of evolving nutrient limitation. A steady reserve of energy-rich LD, despite the nutrient-poor settings, may equip cells biophysically to secure immediate competitive advantages (*44*, *45*, *51*, *52*), while preserving essential molecules for the maintenance (or modification) of behavioural and physiological traits over much longer timescales.

As nutrient limitation persists, isolated LDs merge to form larger droplets, reducing their total count per cell. The enhancement of the effective lipid volume (Figure 1B, Methods) is accompanied by directed cytoplasmic translocation of the LDs away from the fore-end of the cell, ultimately accumulating below the cell nucleus. Inset Figure 1B (multichannel fluorescent image) captures the position of the LD, relative to the cell nucleus (*N*), and the propelling flagellum (*F*). Since the position of the nucleus (*C*_*N*_, Figures M1) coincides with the center of buoyancy, *C*_*B*_, or the geometric center of the cell in an LD-free state, the center of gravity and the geometric center overlap. The center of gravity, obatained by weighted average of the lipid, nucleus and cytoplasmic areas in the cell. Using *C*_*B*_ as the center of wind rose plots (Figures 1C-E), we track the radial and angular coordinates of the cytoplasmic LDs over time (*C*_*L*_). During the early exponential stage (Figure 1C), the LDs are miniscule in size (*V*_*LD*_ < 5 μm^3^), distributed unspecifically between both the top-half and the bottom-half of the cytoplasm. Due to their small size, and close proximity to the geometric center (short *L*_*L*_, Figure M1), LDs do not alter the orientational stability of cells (Methods, Figures 3 and 4). However, during the stationary stage, higher LD volumes (5 μm^3^ < *V*_*LD*_ < 20 μm^3^), combined with predominant localization below the cell nucleus, modify the center of gravity of the cells (Figure 1D). Larger LDs, placed further away from the geometric center of the cell, increase the offset distance of the LDs, *L*_*L*_, thereby altering the separation between the geometric center and the effective center of gravity. A shift in the offset length generates rotational moments (Figure 4), biomechanically modifying the orientational stability of swimming cells (*48*). Over the course of the stationary stage, progressive accumulation of LDs, with specific gravity (~0.9) lower than the surrounding cytosol (~1.05), relocates cells’ center of gravity above the geometric center, *i.e.,* toward the upper half of the cell (Figures 1C-E, Figure M1). The new coordinates of the center of gravity is underpinned by the LD size and the offset length. Since LDs are lighter than the cytosol their intracellular mobility along the gravity vector (*i.e.*, against the direction of phytoplankton migration, Figure 3A, B) suggests that cytoplasmic translocation is not a passive gravity-mediated process, but rather an active process. During the late stationary stage, LDs do not translocate further (Figures 1D, E), however their growth relative to the cell size, generates stronger rotational moments, with far-reaching impact on the migratory properties (Figures 3 and 4).

A similar trend is observed for S-2, however with different LD growth and translocation dynamics (Figure 1F-J). Compared to S-1, the normalized lipid size, *V*_*LD*_, shows an increasing trend even into late stationary stages (Figure 1G), due to increasing lipid volumes (inset, Figure 1G), and relatively stable cell sizes. Akin to S-1, the LDs translocate progressively below the nucleus, as captured in the wind rose plots in Figures 1H-J. It might be worthwhile to mention here that the nucleus of S-2 is located above the geometric center of the cell (*48*), thus making the center of gravity slightly higher than the geometric center of the cell (top heavy) in the LD-free state. As LDs grow and translocate below the cell nucleus, the effective center of gravity moves lower, reducing its separation from *C*_*B*_. In summary, both S-1 and S-2 elicit growth and translocation of LDs, however maintaining strain-specific differences in the LD growth rates (0.0895 μm^3^/h for S-1 vs 0.044 μm^3^/h for S-2), intracellular translocation speeds (at 0.005 μm/h for S-1 vs 0.0033 μm/h for S-2), and cell size modifications, allowing for selective regulation of swimming characteristics in a strain-specific manner.

Reincorporation of nutrients into the depleted cell cultures reconfigures the LD growth and translocation dynamics. Within ~10 h of nutrient reincorporation, LDs shrink in size, and thereafter break down into droplets of smaller sizes. The lysed droplets now translocate in the reverse direction, moving toward the fore-end of the cell, ultimately disappearing within the course of the population doubling time (Figures 2). The cell size does not alter during lipolysis (Figure 2A), while the freshly lysed smaller droplets continue dispersing across the cytosol (false color fluorescent micrographs, Figure 2A). Quantifying the LD size, we find that the net lipid volume per cell drops significantly between 10 h and 24 h (Figure 2B) (stat given in the caption already), finally reducing to ~5 μm^3^ within ~35 h post reincorporation, *i.e*., within the doubling time of the population (specific exponential growth rates for S-1 and S-2 are 0.66/day and 0.4/day respectively, in agreement with previous studies (*48*). With ~50% of the cells retaining *V*_*LD*_ below 5 μm^3^ after reincorporation (Figure 2E), at a population level, the S-1 elicits a higher diversity of cells having lipids. The reversed translocation of LDs toward the fore-half of the cell (Figures 2C-E), consequently shifts the center of gravity back at the geometric center of the cell. The rate of LD lipolysis, tunable by the concentration of the nutrients reincorporated, is considerably faster than that of lipogenesis (Methods). Lipolysis in S-2 is more rapid, with significant reduction of the cellular lipid within the first 6 h - 10 h of nutrient reincorporation (Figures 2F, G). Within ~35 h, the LDs reduce by 97% of the initial *V*_*LD*_ (*V*_*LD*_ < 5 μm^3^), within 80% of all the cells chosen randomly (Figures 2H-J), thus eliciting a uniform, low diversity population compared the S-1. The reconfigurability of the LDs–both in size and spatial localization–confirm that cytoplasmic translocation of LDs is an active process driven by the nutrient availability in the local environment. While strain-specific differences are noted (and contextualized in following sections), for instance lipolysis in S-1 is slower than in S-2, and that a fraction of the S-1 population retains LDs even after reincorporation, in contrast to S-2, overall both strains manifest an active nutrient-controlled growth and spatio-temporal dynamics of intracellular lipid droplets.

**Figure 2.**
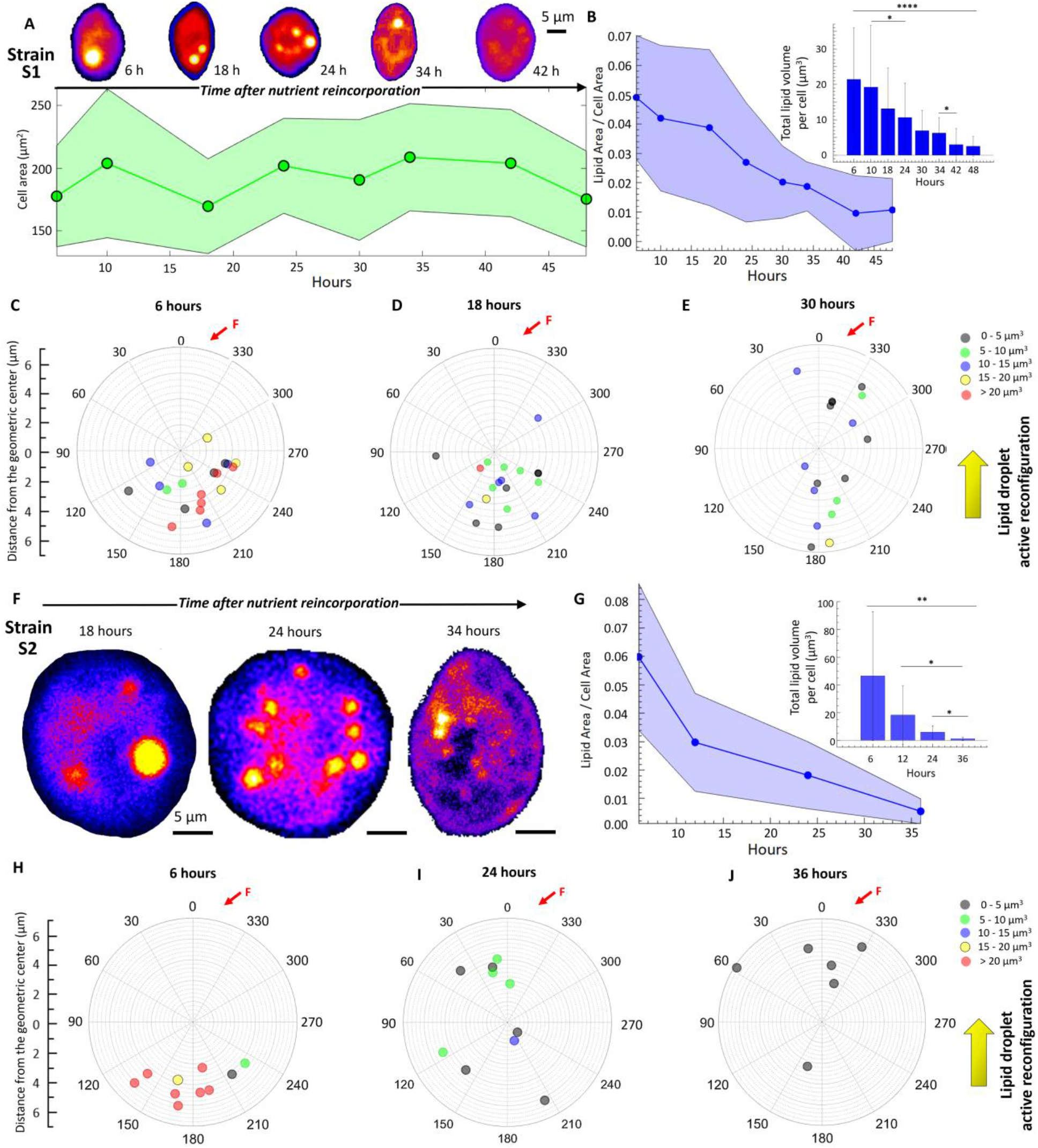
Active reconfiguration of LDs upon nutrient reincorporation. **(A)** Rapid lipolysis of cytoplasmic LDs (bright spots) in S-1 upon reincorporation of nutrients in the cell culture. The cell area remains nearly constant across the observed time window. Shown here using false color fluorescent micrographs, within 18 h of reincorporation, large LDs break down into smaller droplets while reducing in size, before disappearing at *t* = 48 h. The lysed and shrinking LDs actively translocate from the aft-to-fore direction (bottom to top), in opposite sense of the growing LDs under nutrient-limited conditions. **(B)** The index of lipid size, *I*_*LD*_ (*I*_*LD*_ = total lipid area/cell area), reduces from 0.049 ± 0.021 just prior to reincorporation, to 0.01 ± 0.01 at *t* = 48 h. Inset shows the reduction in the total LD volume per cell, *V*_*LD*_, over time after nutrient addition. One-way ANOVA between 10 h, 18 h and 24 h, t-test between 34 h and 42 h, p < 0.001; asterisks indicate statistically significant difference. **(C-E)** Spatio-temporal coordinates of the effective LD in individual S-1 cells after nutrient reincorporation, measured relative to the geometric center (*C*_*B*_). Effective LD size in each cell reduces, and the intracellular positions are shown for *t* = 6 h **(C)**, 18 h **(D)**, and 30 h **(E)** respectively. An active translocation of LDs can be seen from the bottom to the top of the cell (indicated by the flagellar position, *F*). **(F)** False color micrographs of S-2 reveal lipolysis, and LD size reduction due to nutrient reincorporation, alongside active translocation of the LDs in the aft-to-fore direction. **(G)** *I*_*LD*_, the total lipid area/cell area, reduces from 0.06 ± 0.025 prior to reincorporation, to 0.005 ± 0.004 at *t* = 36 h. Inset shows the reduction in the total LD volume per cell, *V*_*LD*_, over time after nutrient addition. One-way ANOVA between 12 h, 24 h and 36 h, t-test between 24 h and 36 h, p < 0.001; asterisks indicate statistically significant difference. **(H-J)** Coordinates of effective LDs relative to the geometric center at *t* = 6 hours **(H)**, 24 hours **(I)**, 36 hours **(J)** respectively capture the active reconfiguration of the LDs due to nutrient reincorporation.

**Figure 3.**
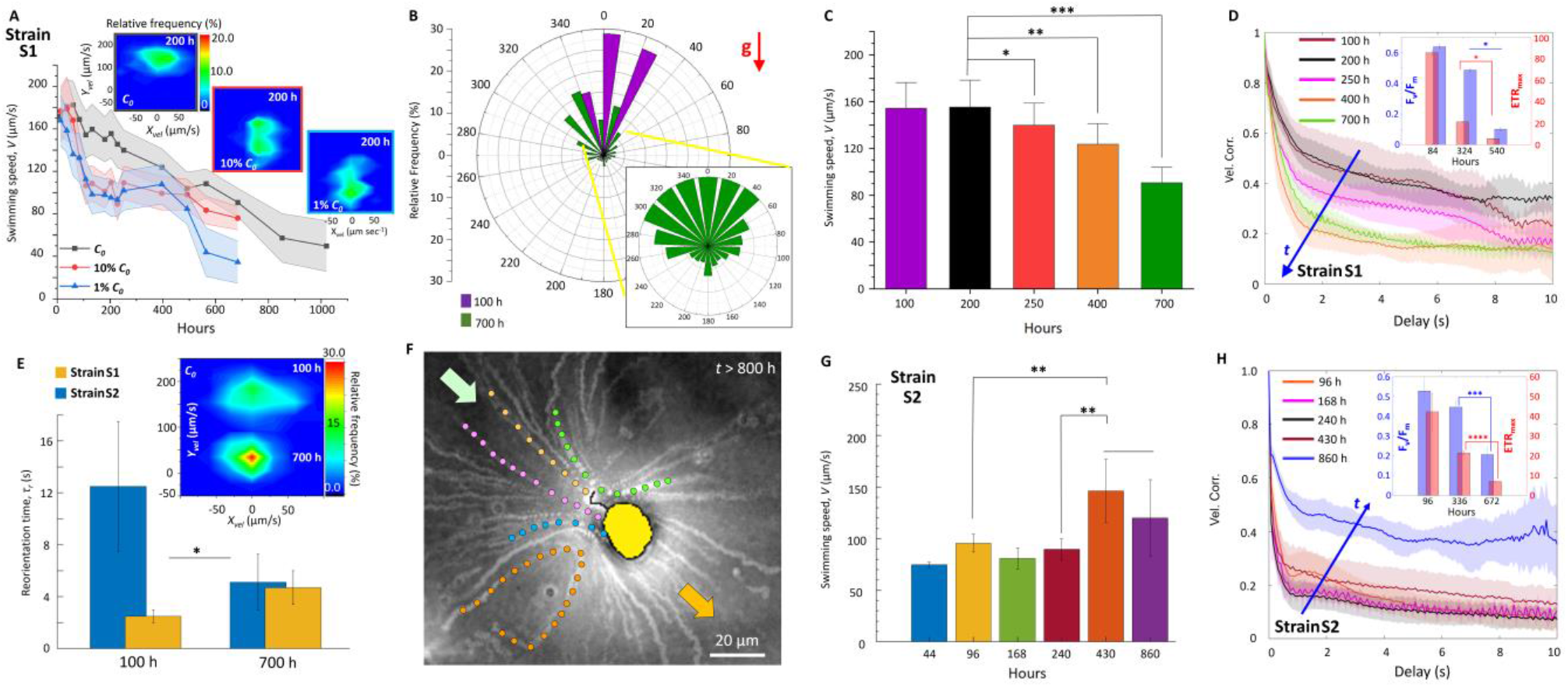
Reconfigurable lipid droplets govern migration of nutrient-limited phytoplankton. **(A)** Evolution of the S-1 swimming speeds spanning the age of the cell culture, with distinct starting nutrient concentrations of the inoculum: Control (*C_0_*, black), 10% *C_0_* (red), and 1% *C_0_* (blue). The corresponding shaded regions depict the statistical confidence (SD). S-1 exhibits maximum swimming speed during the early exponential growth stage (for each starting concentration), and reduces progressively as the culture reaches late exponential and stationary stages. The starting nutrient concentration in the cell inoculum determines the rate at which the cell cultures ages (time taken to reach the late stationary stage). Inset (left to right): Joint velocity distributions corresponding to 200 h since inoculation for starting concentrations *C_0_*, 10% *C_0_*, and 1% *C_0_* reveal emergence of sub-populations with distinct vertical migration behavior. **(B)** Windrose plot captures the swimming directionality of S-1 at *t* = 100 h (purple) and *t* = 700 h from the start of the control inoculum (*C_0_*). The younger cell population exhibits strong anisotropic motility along the vertical direction (against the gravity, **g**), whereas the older cell population loses swimming anisotropy, as revealed by the angular spread of the swimming directions. The zoomed-in view of the windrose center captures the emergent migration of the older cell population along the gravity direction. **(C)** Variation of the swimming speed, *V*, of S-1 population spanning 100 h – 700 h of the control inoculum (*C_0_*). *V* decreases significantly as the duration under nutrient-depletion is extended (*t* > 200 h). The significant differences (one-way ANOVA, p < 0.001, post-hoc Tukey’s honest significant difference) between the exponential (*t* = 100 h and 200 h), late exponential (*t* = 250 h) and stationary (*t* = 400 h and 700 h) growth stages are observed. The bar plots represent the mean swimming speed ± s.d., one asterisk indicates statistical significant difference to 100 h and 200 h; at 400 h, two asterisks indicate statistical significant difference relative to swimming speeds at 100 h, 200 h, and 250 h. At 700 h, three asterisks indicate statistical significant difference to 100 h, 200 h, 250 h and 400 h. **(D)** S-1 velocity correlations across different time points of the population growth. The loss in the velocity correlation signals a shift from ballistic (*t* = 100 h) to a diffusive (*t* = 700 h) motility regime over the course of the population growth indicated by the arrow head. The inset shows significant reduction of the photosynthetic performance concomitantly, measured in terms of the photosynthetic efficiency, *F*_*v*_/*F*_*m*_, and the maximum electron transfer rate, *ETR_max_*. t-test between 324 h and 540 h for both *F*_*v*_/*F*_*m*_, and *ETR_max_* p < 0.001; asterisks indicate statistically significant difference. **(E)** Orientational stability of S-1 population, measured as the reorientation time, *τ_r_*, at *t* = 100 h and at *t* = 700 h. Higher orientational stability corresponds to a lower *τ*_*r*_, i.e. faster reorientation back to the stable swimming direction after the population experienced an orientational perturbation (see Methods). The bar plot represents mean ± s.d, and the asterisk indicates statistical significance between the reorientation time of exponential and stationary stages (two sample t-test, p < 0.00). Inset: Joint velocity distribution captures the relative shift in the vertical velocity component, *Y_vel_*, from *t* = 100 h to *t* = 700 h in S-1 population. The corresponding horizontal velocity component, *X_vel_*, shows a peak frequency around ~0 μm sec-1 for both the time points. **(F)** Time averaged micrograph obtained from a movie of a phytoplankton generating feeding current under nutrient depletion (*t* > 800 h). The bright streaks indicate trajectories of bacteria and passive particles within the feeding current, a selection of which is shown with different hues. The loss of vertical motility is accompanied by the generation of feeding current, suggesting a shift from phototrophic to phagotrophic foraging mode under nutrient limitation. **(G)** Swimming speed of S-2 population spanning 44 h – 430 h of the control cell inoculum. In contrast to S-1, S-2 migrate faster against gravity as the duration under nutrient-depletion is extended (*t* > 240 h). One-way ANOVA between 96 h, 168 h, 240 h and 430 h, t-test between 240 h and 430 h, p < 0.001; asterisks indicate statistically significant difference. **(H)** S-2 velocity correlation shows an increase with the age of the cell culture, i.e., under extended duration of exposure to nutrient limitation. The increase in velocity correlation with age, especially a significant increment during the late stationary stage, signals an altered swimming behavior toward ballistic regime (*t* = 430 h). Similar to S-1, a significant reduction of the photosynthetic performance accompanies the altered motility (inset), measured as *F*_*v*_/*F*_*m*_, and the maximum electron transfer rate, *ETR_max_*. t-test between 336 h and 672 h for both *F*_*v*_/*F*_*m*_, and *ETR_max_* p < 0.001; asterisks indicate statistically significant difference.

**Figure 4.**
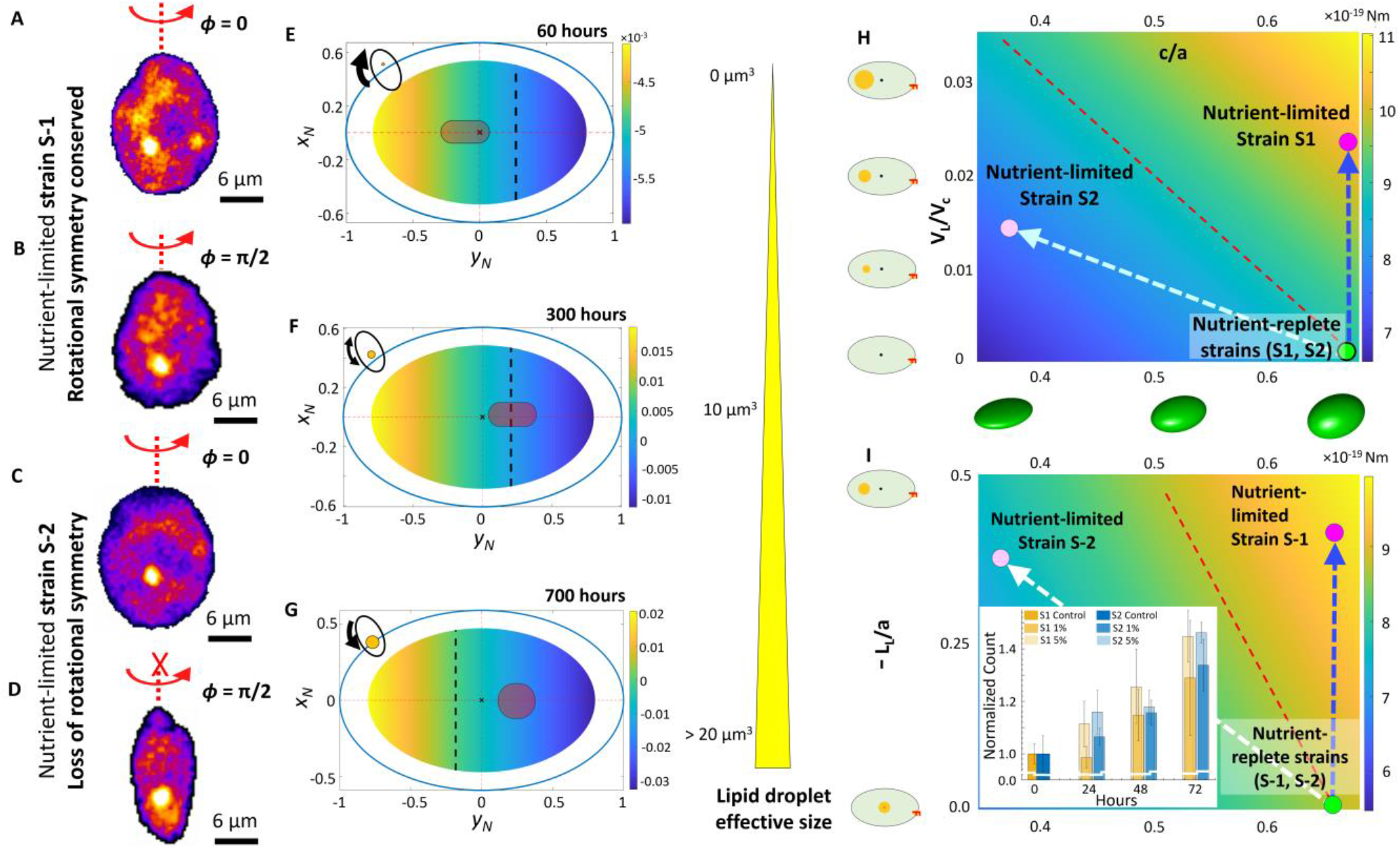
A synergistic interplay of active LD translocation and morphological pliability governs strain-specific adaptive migration under nutrient limitation. **(A-D)** The variation in cell shape in nutrient-replete and nutrient-depleted conditions for S-1 (panel A and B) and S-2 (panels C and D). Cells of S-1 population retain axisymmetric morphology during the course of nutrient depletion, while for the S-2 cells, the ratio of *c* and *b* reduces from ~1 to 0.53 ± 0.13. **(E-G)** Phase plots delineating the orientational stability of a swimming cell as the position of cytoplasmic LDs change (based on experimental data presented in Figures 1C-E), mapped against the corresponding reorientation speed from an unstable to a stable orientation (Methods). The phase plots are obtained from experimental data, accounting for the morphology, nucleus size and posiiton relative to the geomtric center, effective lipid size and position, and swimming velocity corresponding to the respective growth stages. The colorbar represent the magnitude of the angular speed about its geometric center. The phase diagram is obtained by varying all possible locations over which LDs may reside (changing *φ*_*L*_ and *L*_*L*_). For the angular speed estimation, the cytoplasm density was taken to be 1050 kgm^−3^ (*16*), nucleus density as 1300 kgm^−3^ (*17*) and lipid density as ~900 kgm^−3^ (*18*). The shaded portion in the above plots delineates the possible positions a lipid droplet may actually be observed (plotted using the mean and the variance from experimental observations, Figures 1C-E). **(H-I)** Phase plot of active torque required by a cell to reorient itself as an upward swimmer (negative gravitactis) for the combination of the ratio of *a* and *c* and (H) ratio of the lipid and the cell volume, and (I) ratio of the lipid position from the cell geometric center and *a*. The green circle shows nutrient-replete cells while magenta circles show nutrient-depleted cells (magenta indicates S-1, light magenta S-2). The dashed red line shows the iso-active torque line. Active torque required by a cell increases with nutrient depletion (when lipid droplets increase in size and translocate opposite to the position of the flagellum). However, the change in the active torque varies across strains and gives a strain-specific behavioral shift. Inset panel (I) shows the daily variation in the normalized cell count of nutrient depleted cells upon 1% *C_0_* and 5% *C_0_* inoculation of f/2-Si medium (added every 24 hours, Methods). S-1 which is known to have a higher growth rate (> 0.65/day), in agreement with (*57*) compared to S-2 (0.4/day, (*48*)) shows a diminished recovery rate at very low availability of nutrient concentration.

The migratory behavior under nutrient limitation is strain-specific, emerging concurrently with commensurate alterations in physiological and trophic traits (Figures 3). By mapping the swimming properties with the intracellular lipid attributes at corresponding time points, we uncover mechanistic links between the LD dynamics and the ballisticity and orientational stability of nutrient-limited cells (Figure 4). For S-1 (Figure 3A-F), the swimming speed, 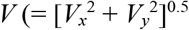, *x* and *y* indicate directions perpendicular and parallel to the gravity vector, ***g***, respectively), quantified within custom-built millifluidic chambers (Figure M2, Methods), is progressively arrested as nutrient concentration drops, triggering emergence of a distinct sub-population with reduced motility (Figure 3A-C). The relative proportion of low motility swimmers increases as the population enters late stationary stage, captured in the joint velocity distribution (at *t* = 100 h and 700 h, Figure 3E, inset). The rapidity with which a nutrient-depleted population switches to low motility regime depends on the initial nutrient availability (Figure 3A, inset): in the control case (initial concentration *C_0_*), the sub-population appears at 200 h < *t* < 250 h (Figure 3C), while for the populations inoculated with 10% *C_0_* and 1% *C_0_*, low motility sub-population appeared within 100 h < *t* < 150 h (Figure 3A). The drop in the swimming speed, specifically, occurs due to significant reduction of the vertical swimming speed, varying from 140.34 ± 37 μm/s during the exponential stage to 19.83 ± 24.47 μm/s in late stationary stage, with the horizontal speed reducing from 62.96 ± 20.8 μm/s in exponential to 30.9 ± 64 μm/s in stationary stage (Figures 3B, 3E (inset)). The broadening of the distribution of swimming direction (Figure 3B), indicates a shift in the migratory strategy from an anisotropic vertical swimming to enhanced horizontal movement under nutrient stress (Figures 3B), in line with migratory adjustments due to other stressors, including temperature (*33*), light and turbulence (*48*, *49*). Under nutrient limitation, the index of anisotropy, *I*_*A*_ (= |*V*_*y*_ / *V*_*x*_ |), drops from 2.1 to 1.32 as the population enters stationary growth stage, synchronizing with the significant increment of the cytoplasmic lipid volume (Figure 1B). Since *I*_*A*_ drops more rapidly in cell cultures inoculated with low initial nutrient concentrations (10% and 1% of *C_0_*, Figure 3A), an inverse relation between the vertical motility and the intracellular LD content is concluded.

The shift in the swimming traits accompanies commensurate modification of photophysiology and trophic properties of *H. akashiwo* strains. The population-scale swimming trajectories allow us to extract the mean squared displacement (MSD) values, signifying the loss of directional swimming and emergence of isotropic swimming traits (Methods). For S-1, the decrease in the velocity correlation over the physiological growth stages indicates a progressive shift from ballistic to diffusive swimming trait (Figure 3D, between *t* = 100 h to *t* = 700 h). Concomittantly, the reorientation time, *τ_r_*, the characteristic time cells take to rotate back to equilibrium orientation once perturbed from it (*48*, *49*), increases from 2.5 ± 0.5 s (*t* = 72 hours) to 4.7 ± 1.3 s (*t* = 700 hours), confirming reduced orientational stability of the nutrient-limited cells (Figures 3E, Methods). Overall, the alteration of the migratory properties indicate a concerted shift in the cells’ ability to span the horizontal space under nutrient limitation, suppressing the vertical migration. Together with the observations of enhanced production of reactive oxygen species (ROS), an indicator of physiological stress (*49*), and the reduction of the photosynthetic performance (*49*, *53*), quantified in terms of the photosynthetic efficiency, *F*_*v*_/*F*_*m*_, and the maximum electron transfer rate, *ETR*_max_ (inset Figure 3D, Methods), we infer that the migratory shift goes hand-in-hand with modification of the physiological traits. Significant drop in both *F*_*v*_/*F*_*m*_, and *ETR*_max_ was recorded between 324 h < *t* < 540 h (the t-test, p < 0.001; asterisks indicate statistically significant difference), which alongside enhanced non-photochemical quenching (NPQ) and altered migratory behavior, provide hallmarks of an immenent trophic modification. A comparison of the timelines reveals that the reduction in *F*_*v*_/*F*_*m*_ follows LD biogenesis and translocation (Figure 1A, B), suggesting that the downregulation of the photosynthetic machinery is correlated with the time of the LD-governed migratory shifts. While the downregulation of photophysiology may serve as an adaptive strategy to conserve energy under nutrient constraints (*54*), it comes at the cost of reduced growth and cell division, thus necessitating alternative trophic strategies for survival. Further prolongation of the nutrient limitation (*t* > 800 h), indeed reveals a switch from photo-to phagotrophic mode of resource acquisition in S-1, evidenced by the generation of feeding currents (Figure 3F). The bright streaks indicate the trajectories of bacteria and passive particles within the feeding current, a selection of which is shown with different hues. The suppression of vertical motility, together with reduced photosynthetic performance and generation of feeding currents equip cells to shift from photo-to phagotrophy under nutrient limitation, enabling them to execute basic metabolic functions till inorganic nutrients reappear (*54–56*).

The shifts in motility and trophic strategies are reversed upon spiking the nutrient-depleted cell cultures with low concentration of fresh nutrient. The uptake of low concentrations of freshly available NO_3_^−^ and PO_4_^3–^ allow nutrient-limited cells to ameliorate physiological stress, recover photophysiology, and enhance vertical migration. After 24 hours of incubation with nutrients at a concentration ~ ⅛ *C_0_*, S-1 showed up to 28% recovery in the photosynthetic efficiency (*F*_*v*_/*F*_*m*_), along with partial recovery of the ballisticity and orientational stability of the cells. The concerted restoration and motility and photosynthetic traits accompany reduction of ROS levels, suggesting that the population is now primed to execute vertical migration toward light-rich upper regions in the water column. The recovery of physiological and behavioral traits, even with a tiny spike in the nutrient concentration, provides motile phytoplankton the crucial ability to execute vertical migration, and perform photosynthesis to increase fitness.

Despite qualitative similarities in the LD biogenesis and cytoplasmic translocation (Figures 1 and 2) of strains S-1 and S-2, their emerging migratory traits differ considerably. Unlike S-1, cells in S-2 do not show signatures of a trophic switch to phagotrophy. Both swimming speed and orientational stability increase as the nutrients turn limiting for S-2 (Figures 3G, H), showing an inverse relation of motility with the nutrient availabity. As the population enters stationary stage, the swimming speed increases from 89.64 ± 10.25 to 146.41 ± 30.47 μm/s over the course of 240 h < *t* < 430 h, due primarily to the higher vertical speeds against gravity (Figures 3G). The velocity correlations increase as the nutrient depletion persists, eliciting a significant rise as the population enters late stationary stage (from *t* = 430 h to 860 h), signalling a behavioral shift from diffusive to ballistic regime (Figures 3H). This, accompanied by significant reduction of the reorientation time from 12.5 ± 5 s to 5.11 ± 2.16 s (*i.e.*, increased orientational stability, Figures 3E), represents a migratory switch that allows the S-2 population to swim effectively against gravity and distribute across the light-rich upper layers of the water column. This is consistent with the higher light adaptivity of S-2, relative to S-1, reflected by the respective NPQ values: for S-2, the NPQ shows anamolous change (Figure 4) and throughout the growth stages, remains consistently lower than that of S-1. Though the photosynthetic efficiency and the electron transfer rate are adversely impacted due to the nutrient limitation (t-test between 336 h and 672 h for both *F*_*v*_/*F*_*m*_, and *ETR*_*max*_ p < 0.001; asterisks indicate statistically significant difference, Figure 3H, inset), S-2 performs relatively better with their values remain lower than those in S-1. Taken together, our results indicate that S-1 upregulates the light-protection mechanism to dissipate excess light energy, whereas S-2 is better light-adapted and attempts to maintain their photosynthetic performance while waiting for chanced encounters with ephemeral nutrient molecules. The alteration of migratory properties act synergistically with the photophysiological attributes (high NPQ and reduced photosynthetic performance), enabling S-2 to position within the upper photic layer, and maximizing the chances of fitness enhancement through chanced encounters with ephemeral nutrient patches (*20*, *35*).

Reconfigurable LDs actively govern the vertical migration of nutrient-limited motile phytoplankton. Alongside LD growth and cytoplasmic translocation, the cells–in a strain-specific manner–undergo morphological changes, whereby S-1 cells become smaller, while S-2 cells become flatter (platelets) losing the rotational symmetry about the cell’s long axis (Figures 4A-D). We develop a computational model for cell mechanics to delineate how LD size and intracellular location, in conjunction with morphological traits, shape the swimming properties of motile phytoplankton. For S-1, accumulation of LDs below the cell nucleus, mediated by directed translocation (Figure 1), shifts the center of gravity above the geometric center, thus rendering the cell slightly *top-heavy* (*48*) relative to the LD-free condition under nutrient replete conditions. Top-heavy cells take longer to orient back to their stable swimming direction when perturbed therefrom, resulting in longer reorientation timescales, *τ*_*r*_ (Figure 3E, Methods) and lower angular reorientation speeds (Figures 3E-G). Equivalently, the active torque (and hence the rotational kinetic energy) required to keep the cells oriented along their initial swimming direction increases as the LDs grow in size (*V*_*L*_, Figure 3H), and the distance from the geometric center increases (*L*_*L*_, Figure 3I). Taken together, as the LDs grow and accumulate below the nucleus, cells become progressively gain orientational stability in opposite direction, thereby shifting the swimming trait from negative to positive gravitaxis.

When a stable upward swimming cell under nutrient replete conditions (Figures 1 and 3) is perturbed to a horizontal configuration (θ = 90^0^ with the cell long axis perpendicular to the gravity vector, Figures 4E-G), it tries to regain the stable swimming direction by reorienting back. The stable swimming direction (upward and downward are denoted by +ve and –ve angular speeds respectively), and the rapidity of reorientation captured by the magnitude of the angular velocity, are underpinned by the intracellular organelle distribution and propulsion forces, Cells in the exponential growth stage reorient upward (negative gravitaxis) with positive angular velocity, owing to the size and location of the LDs lying within the shaded portion (grey hue based on experimental data, Figure 4E). This confirms the high ballisticity and low reorientation times observed during the exponential growth stage (Figures 3D, E). During the late exponential stage (*t* = 300 h, Figure 4F), cells become neutrally stable, with similar probability of turning stable or unstable in the direction of gravity vector. Owing to the low LD volume, and localization in the region of vanishing angular speed (grey hue close to the dashed line, Figure 4F), ballisticity of swimming reduces, with increase in the reorientation time (*i.e*., the angular speed changes to –ve sign with increased magnitude, as shown by the color bar). This is validated by the reduced swimming speed and ballisticity as the population transitions to stationary stage (Figures 3C, D). Finally, cells from the late stationary stage (*t* = 700 h, Figure 1E), are strongly stable in the downward direction (positively gravitactic), due to the large LD size and conspicuous localization below the nucleus, eliciting a strong reorientation speed ~ −0.01 s^−1^ (Figure 4G). Since the shaded portion falls in the region of the unstable angular speed (to the right of the dashed line), cells of S-1 would require sufficient active torque in the other direction so as to become stable upward swimmers (Figures 3H, I).

Our mechanistic model, in contradiction to our experimental observations (Figures 3G, H), predicts that the cells from S-2 being top-heavy under nutrient replete conditions (*48*), would turn highly stable down swimmers (positively gravitactic) as LDs progressively appear. We reconcile this apparent discrepancy by accounting for the concomitant morphological changes in S-2 under nutrient limitation. While S-1 preserves the rotational symmetry under nutrient depletion, thus maintaining the ratio between the lengths of the semi-minor (*b*) and semi-major (*a*) axes, *b/a* ~0.65 at 60 h and ~0.6 at 696 h (Figures 4A, B), cells of the S-2 turn into platelet morphology, thus losing their axial symmetry in face of nutrient depletion (Figure 4C, D). The platelet morphology is quantified by the degree of flatness, (*c/a*)*^−1^*, where 2*c* is the maximum dimension orthogonal to both *a* and *b* (lower the value of *c/a*, flatter is the cell). The value of *c/a* reduces from ~0.7 to ~0.31, while *b/a* varies from ~0.7 at 96 h (*48*) to ~0.6 at 860 h, resulting in lower active reorientational torque to maintain negative gravitaxis during the late stationary stage relative to its early exponential counterpart. By mapping *c/a* against growing lipid volume (Figure 3H) and the cytoplasmic position (Figure 3I), we compute the active torque required to maintain negative gravitaxis (phase plot in Figure 4H–I). The active torque required for S-2 cells to assume a stable configuration attenuates as nutrient depletion persists, emerging due to an interplay between the cell flatness and the combined effect of LD size and their localization within the cytoplasm. The reduction in the active torque follows directly from the corresponding drop in the rotational viscous resistance (~30%) experienced by platelet-shaped cells under prolonged nutrient limitation. In summary, LDs, in synergy with strain-specific morphological traits, enable cells to differentially fine tune migratory properties in face of nutrient limitation. Herein, the rotational symmetry in cell shape about the cell’s long axis plays a key role in regulating the vertical migration. This morphological pliability allows S-2 to harness the spatio-temporal dynamics of cytoplasmic LDs to enhance vertical motility, in contrast to S-1. This differential, strain-specific LD-governed migratory trait conforms to the measured physiological changes (ROS, photosynthesis and trophic strategy) under nutrient limitation and upon reincorporation of nutrients (Figure 4I inset).

## Discussion

Under nutrient limitation, motile phytoplankton species resort to two distinct strategies, which taken together, indicate their ability to maximize resource acquisition and fitness. Strain S-1 (CCMP3107) reduces vertical motility, and switches to phagotrophy, while strain S-2 (CCMP452) enhances vertical migration, switching to stronger gravitatic behavior aided by the morphological pliability. At the level of single cells, the strain-specific contrasting strategies emerge due to the active biomechanical control exerted by the spatio-temporal dynamics of cytoplasmic LDs on the swimming properties. By spanning different stages of growth in the cells cultures as a proxy of ecologically relevant nutrient concentrations, we quantify the behavioral and physiological traits and underscore their interrelations as coupled traits which enable species to enhance fitness under nutrient limitations. Cells of S-1 require increasing levels of active torque (hence energetically expensive, particularly in face of nutrient limitation) to sustain negative gravitaxis, reflected by the depletion-induced increase in the reorientation time. For the S-2, the active torque required to maintain vertical motility is reduced (so is the reorientation time), due to the reduction of the viscous drag. Leveraging the experimental and computational data, we present a dissipative power budget, accounting for the gain in the gravitational energy and viscous losses incurred by the swimming cells per unit time. The power dissipated reduces for both strains, thereby reducing the active energy input by the cells to sustain motility, however they do so distinctly. S-1 suppresses motility and reduces cell size to avoid the viscous dissipation under nutrient limitation, whereas S-2 is able to sustain, and even enhance vertical motility, benefitting from a lower rotational and translational viscous drags, and hence energy dissipation under prolonged limited settings. For S-1, the energy dissipation drops from 3.75 fW to 1.18 fW as the population transitions from exponential to stationary growth stage; while for S-2, the dissipation decreases from 6.31 fW to 3.24 fW, as the cell transforms into platelet morphology during the late stationary stage. These biomechanical changes and energetic insights are backed up by the alteration of physiological traits, specifically shift in the trophic mode (photo-to phagotrophy in S-1, Figure 3F) and accompanying strain-specific photophysiological changes (Figures 3D and H), ultimately enabling the strains to maximize fitness by repositioning across distinct vertical depths in natural environments.

The dynamic tuning of active torque and dissipative power under nutrient limitation suggests that motile phytoplankton may have exquisite decision making abilities, complementing intrinsic mechanisms (*58*). While S-1 cells reduce dissipative power progressively as nutrition depletion persists, S-2 first increases dissipative losses by enhancing active torques (from early exponential to early stationary growth stage), thereafter reducing the active torque and dissipation aided favorably by the morphological changes over longer periods of nutrient depletion. The ability to arrive at a physiologically-amenable behavioral trait allows motile phytoplankton to conserve energy dynamically as nutrient landscapes vary. Maintaining vertical motility even under nutrient limitations allows S-2 cells to enhance fitness through chanced encounters with ephemeral molecules as and when they are available (*9*, *35*). We confirm this by comparing the relative growth rates of nutrient-starved populations of S-1 and S-2 upon reincorporation of small amounts of fresh nutrients (1% and 5% *C_0_*, Methods). S-1 grew slower relative to S-2, although under nutrient-replete conditions S-1 shows higher growth rates, suggesting that the strain S-2 has a higher nutrient affinity, *i.e.*, better ability to utilize miniscule amounts of nutrients as and when they are available (inset, Figure 4I) for the first 48 hours, after which S-1 cells match up the growth rate of S-2. This is further backed by our measurements showing rapid lipolysis of S-2 cells compared to S-1 (Figure 2) upon nutrient reincorporation. Our data suggest that motile phytoplankton may harness contrasting inter-strain behavioral traits to maintain population viability via distribution of labor and energy resources under nutrient limited conditions (*59*, *60*).

**Figure 5.**
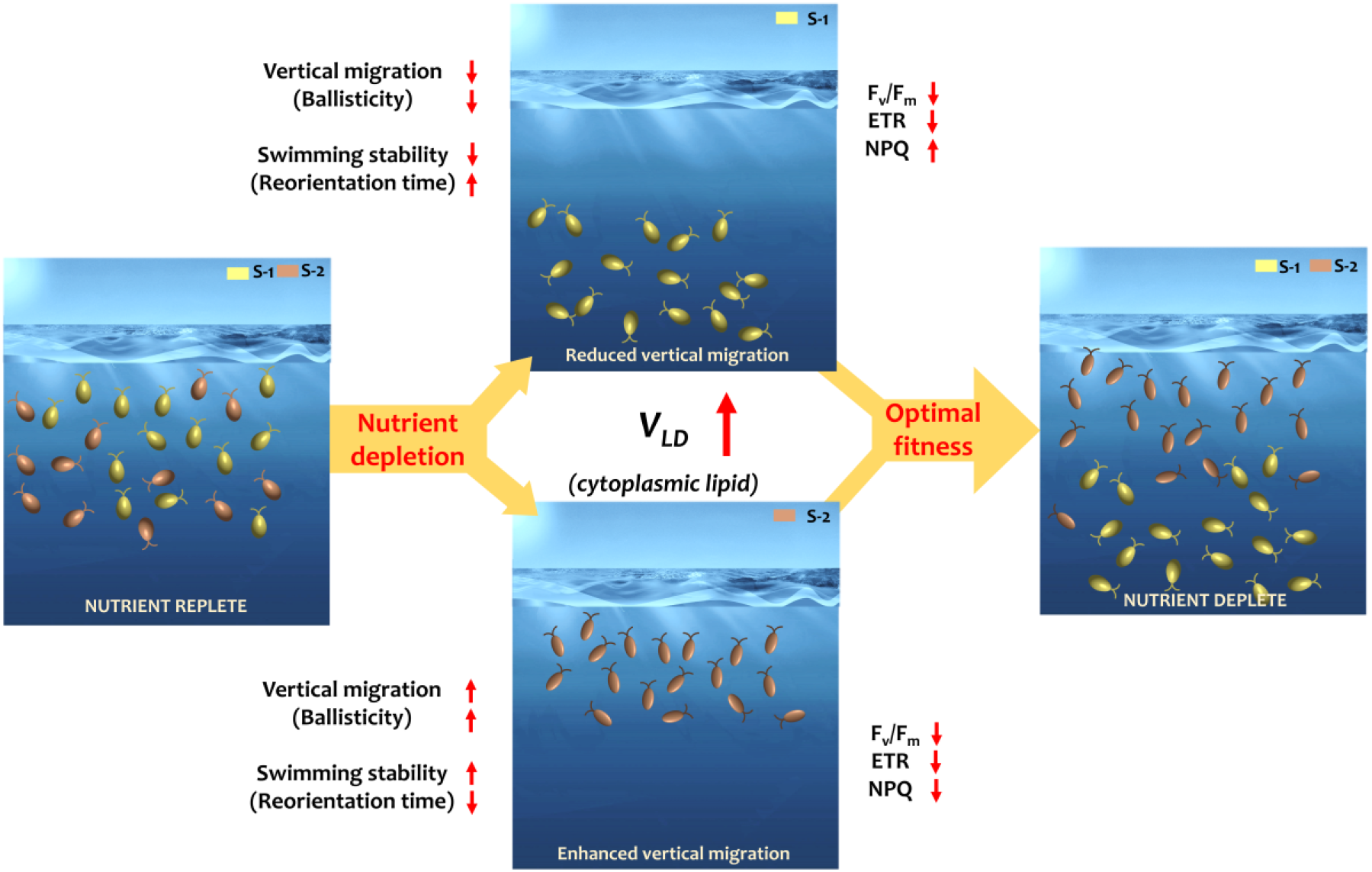
LD-based energy storage and strain-specific migratory shifts act concertedly as coupled traits to enhance fitness of nutrient-limited phytoplankton populations. Naturally co-existing strains of motile phototrophic *H. akashiwo*, S-1 and S-2, execute diel vertical migration under nutrient replete conditions. The phytoplankton population carry out photosynthesis, shuttling between light-rich photic zone during the day time and nutrient-rich depths at night. While S-1 exhibits stronger motility (high ballisticity and low reorienation time), and higher photosynthetic efficiency (higher *F*_*v*_/*F*_*m*_ and *ETR*_*max*_ values) relative to S-2, it is less adapted under high light conditions indicated by the lower non-photochemical quenching (NPQ) parameter. As nutrient limitation sets off, both strains generate cytoplasmic LDs, while distinct behavioral and physiological shifts co-emerge, indicating that the LD-based energy storage and emergent migratory behavior adapt differentially. In nutrient limited settings, generation of intracellular LDs and their directional translocation drive suppression of vertical migration in S-1, while for S-2, significant enhancement in measured. Enhancement of the vertical motility in S-2 is facilitated by the concomitant morphological change, specifically due to the loss of the rotational symmetry as cells turn flatter in shape (platelets), thereby reducing the viscous losses during swimming. The morphological shift allows S-2 cells to conserve energy while maintaining motility in the upper light-rich surface waters, while for S-1 cells, the loss of ballisticity and orientational stability drive them toward diffusive regime in deeper waters. Alongside, the increased NPQ and reduced *F*_*v*_/*F*_*m*_ values of S-1 conform with our observation their trophic switch from photo-to phagotrophic mode of resource acqusition. S-2, on the other hand, increases their ballisticity and motility, thus increasing the residence time in shallow light-rich waters. A relatively lower NPQ shows their enhanced ability to use the incident light toward photochemical processes. Finally, the reduction of the cell size in S-1, and an increase in the overall cell size of S-2 indicate that the two strains diversify their strategies for resource acquisition under limited settings. Under limited concentration of nutrients, small, low motile cells with high surface are to cell volume ratios will have competitive advantage (a strategy potentially adopted by S-1). In contrast, for stable cell nutrient quota, increased size is advantageous (*2*), which may allow S-2 to acquire just enough nutrients, e.g., upon chanced molecular encounters, for growth and division thus enhancing fitness during nutrient limited conditions (inset, Figure 4I). These differential strain-specific responses suggest that motile phytoplankton possess concerted decision-making mechanisms that allow contrasting adaptive strategies for acquiring a limited pool of nutrients, while maximization population-scale fitness. In doing so, phytoplankton redistribute along the vertical column, thus cumulatively covering larger physical space, while minimizing the overlap and potential inter-strain competion for limited resources.

Understanding how phytoplankton adapt and survive the rapidly evolving nutrient landscapes of today’s oceans remains a crucial challenge. Accurate prediction of the cascading biogeochemical implications will rely on mechanistic understanding of co-emerging behavioural and physiological responses, and the interrelations therein (*2*, *61*). Our data suggest that migration and nutrient storage–so far considered indepedent traits (*35*)–co-evolve in a concerted fashion to maximize population fitness, via seemingly contrasting strategies to accommodate strain-specific constraints. The role of lipid droplets in regulating the biomechanics of phytoplankton migration may be conserved evolutionarily among eukaryotic swimmers, thus future efforts could be directed toward understanding the molecular underpinnings of the migration-storage interrelations. In addition to providing new mechanistic insights into the paradoxical diversifcation of species (*62*, *63*), our findings offer a fresh perspective to the co-emerging adaptive traits in phytoplankton under stressful environments, thereby opening up quantitative avenues to assess impacts of multiple stressors (*1*), beyond and in conjunction with evolving nutrient landscapes, which plague marine environments today.

## Acknowledgements

The authors thank Gea Guerriero for supporting with the cell culture facility during initial phase of the project, and Nicolas Tournier for assisting with the fabrication of the millifluidic setup.

## Materials and Methods

### Cell culture

We focus on two different strains of raphidophyte *Heterosigma akashiwo*, namely, CCMP3107 (S-1) and CCMP452 (S-2) for our investigations. For the experiments, cells were cultured in 50 mL sterile glass tubes under a diel light cycle (14 h light: 10 h dark) in f/2 (minus silica, -Si) medium (C_0_), at 22°C. For propagation of the cell cultures, 2 mL of the parent culture was inoculated into 25 mL of fresh medium every two weeks. For propagation of cultures in nutrient-limited environments, f/2 (-Si) medium was appropriately diluted in Artificial Sea Water (ASW, 38 g/L of sea salts in MilliQ water) to reach 10% C_0_ (10x dilution, 10 mL f/2(-Si) and 90 mL ASW) and 1% C_0_ (100x dilution, 10 mL 10x f/2(-Si) and 90 mL ASW) dilutions. *H*. *akashiwo* 3107, 452 cultures used for phenotypic traits quantification experiments were propagated from a 5-7 days old pre-culture (mid-exponential growth stage) to standardize the starting population physiological status. A fixed period of the day (between 9:00 h and 15:00 h) was chosen for the experiments to rule out any possible artifact due to the diurnal migration pattern of phytoplankton species, including *H. akashiwo*.

### Quantification of lipid droplet volume and intracellular effective size localization

To characterize and quantify the biosynthesis and accumulation of lipid droplet (LD), cells were sampled from the culture tube at different time intervals to cover whole exponential and stationary growth stages and stained with neutral lipids fluorescent stain Nile Red (Thermofisher, excitation/emission 552/636 nm). 10 μL of 100 μM Nile Red (in DMSO) were dissolved into 200 μL *H*. *akashiwo* culture supernatant and mixed thoroughly by vortexing the mixture. 200 μL cells were added to the mixed NR stain aliquot (final concentration 2.4 μM) and incubated in the dark at room temperature for 15 minutes. To identify and characterize the accumulation of LDs in single cells, we used phase contrast and fluorescence microscopy (Olympus CKX53 inverted microscope) supplemented with high-resolution color camera (Imaging Source, DFK33UX265). To avoid any photo-toxicity effect of light during cell analysis, excitation LED intensity (552 nm) was kept low and at a maximum value of 5. To extract the *H*. *akashiwo* 3107 cell area and both LD dimension (area and volume) and intracellular location, movies of single-cell were recorded at 16 frames per second for 5 seconds. We quantified the LD effective size assuming individual LDs to be a spherical droplet, and deriving the effective radius of individual LDs from the contour area extracted by thresholding the Images using in-house MATLAB image processing algorithms and Image J. From the individual LDs, the position of their net center of mass in single cells was determined with respect to the cell geometric center.

### Quantification of nucleus radius and intracellular localization

To determine nucleus size and position, *H*. *akashiwo* 3107 cells were sampled from the culture tube at different time intervals representative of lag (60 hours), exponential (300 hours) and stationary (700 hours) growth stages, and stained with Syto9 Green Fluorescent Nucleic Acid Stain (Thermofisher, excitation/emission 480/501 nm). 7.5 μL Syto9 100 μM (in DMSO) were dissolved into 300 μL *H*. *akashiwo* 3107 cell supernatant and mixed thoroughly by vortexing. 300 μL cells were added to the mixed Syto9 stain aliquot (final concentration 1.2 μM) and incubated in the dark at room temperature for 12 minutes. To quantify nucleus characteristics in single-cell *H*. *akashiwo* 3107, we used phase contrast and fluorescence microscopy (Olympus CKX53 inverted microscope) supplemented with high-resolution color camera (Imaging Source, DFK33UX265). To avoid any photo-toxic effect of light during cell imaging, LED excitation intensity (480 nm) was kept low and at a maximum value of 5. To extract *H*. *akashiwo* 3107 nucleus radius and intracellular localization, movies of single-cell were recorded at 16 frames per second for 5 seconds. We derived the effective radius of the nucleus from the contour area extracted by thresholding and Image J image analysis. Relevant data on *H. akashiwo 452* can be found in refs

### Pulse-amplitude modulated chlorophyll fluorometry (PAM) experiment

PAM was used to evaluate and quantify the photophysiological efficiency of *H*. *akashiwo* cells at different time intervals during the exponential and stationary growth stages in diverse nutrient regimes. Multiple Excitation Wavelength Chlorophyll Fluorometer (Multi-Color-PAM; Heinz Walz GmbH, Effeltrich, Germany) was used to quantify the maximum photosynthetic quantum yield (*F*_*v*_/*F*_*m*_), the maximum electron transport rate (*ETR_max_*) and nonphotochemical quenching (NPQ) of *H*. *akashiwo* cells at the population scale. For PAM measurements, culture tube was mixed through gently rotating the tube by 360°. 1200 μL suspensions of plankton cells were sampled from the top 0.5 cm culture and placed into a quartz silica cuvette (Hellma absorption cuvettes, spectral range 200-2500 nm, path length 10mm). We used Multi-Color PAM 3 Win software saturation pulse (SP) and light curve method to quantify *F*_*v*_/*F*_*m*_ and *ETR_max_* at diverse time intervals, and under different nutrient regimes.

### Nutrient-starved phytoplankton biomechanical and physiological response to fresh nutrients

Nutrient-depleted *H*. *akashiwo* cells from the stationary growth stage exhibit characteristic biomechanical and physiological properties (LD accumulation, motility and reduced photophysiology) that enables phytoplankton population to save energy and survive nutrient constraints. Cultures of *H*. *akashiwo* cells from stationary growth stage (700 h) were supplemented with low amount of fresh f/2(-Si) to evaluate and quantify phytoplankton biomechanical and physiological recovery efficiency during a short time interval of 48 hours. 8 mL Late stationary cell cultures were reincorporated with (i) 1 mL C_0_ f/2(-Si), (ii) 1 mL 10% C_0_ f/2(-Si), (iii) control (no nutrient reincorporation). Total nutrients (nitrate and phosphate) concentrations per cell (quantified to stationary cell concentration of 10^5^ cells mL^−1^) were estimated to be: NO_3_^−^(i) ~ 7×10^−7^ μmol cell^−1^, (ii) ~ 7×10^−8^ μmol cell^−1^, (iii) 0 μmol cell^−1^; PO_4_ ^3−^ (i) ~ 4.5×10^−7^ μmol cell^−1^, (ii) ~ 4.5×10^−8^ μmol cell^−1^,(iii) 0μmol cell^−1^. After 24 hours of incubation at 22°C (14 hours light:10 hours dark), *F*_*v*_/*F*_*m*_, *ETR*_*max*_, total LDs volume per cell, motility and ROS were quantified (as described previously) to determine recuperation of *H*. *akashiwo* motility and physiological recovery efficiency.

### Quantification of nutrient concentration

Analysis of nutrients, namely nitrate (NO_3_^−^) and phosphate (PO_4_^3−^) were performed colorimetrically using Prove600 Spectroquant (Merck). NO_3_^−^ was analyzed with Nitrate Cell Test in Seawater (Method: photometric 0.4 – 13.3 mg/l NO_3_^−^Spectroquant®). The corresponding detection limit of NO_3_^−^ concentration measurable in our experiments is 6.45 μM. PO_4_^3−^ was analyzed with Phosphate Cell Test (Method: photometric 0.2 – 15.3 mg/l PO_4_^3−^Spectroquant®, with corresponding detection limit of 2 μM). Experimental values below the respective detection limits is taken as zero.

### Statistical analysis

We performed one-way ANOVA to compare the total lipid droplet volume accumulation in single *H*. *akashiwo* 3107 among different time intervals from exponential to stationary growth stage. We made multiple comparisons using a post-hoc Tukey’s HSD test. Same multiple comparison statistical analysis was conducted to compare phytoplankton speed and photophysiological parameters among different time interval samples representative of the exponential and stationary growth stages. Same statistical analysis was conducted in the *nutrient recovery experiment* to compare photophysiological efficiency, LDs volume, motility and ROS among the still (control, no nutrients) and cells supplemented with fresh f/2(-Si) after 24 hours of incubation.

### MSD Estimation

The proper quantification of ballistic motility is ensured through a rigorous cell-tracking followed by mean squared displacement (MSD) analysis. For the analysis of MSD and the corresponding velocity correlations (insights and details in the main draft), we closely follow the procedures in (*64*). The package @msdanalyzer is modified and used in MATLAB to obtain the MSD curves for 60 hours, 300 hours and 700 hours culture age. The log-log representation of the MSD plots can be fitted with a linear function, that is, *log(<r^2^>)=Γ+αlog(t)* where the exponent α provides the information whether the flow is diffusive (α~1); ballistic motion / active transport / super-diffusive (α~1.5-2) or constrained transport (α<0.9) in nature. To obtain the exponents, we clip the MSD plots until where the phytoplankton cells do not feel the confinement effect.

### Cell mechanics model and phase plot

To understand the cell stability, dead and active torque and cell reorientation timescale (as obtained from experiments), we propose a reduced-order model for the cell mechanics. Here we talk in terms of force and torque balance on the phytoplankton cell and attempt to decipher the effect of lipid formation on the cell stability and rotational kinetics. The focus of the discussion is to identify the relevant forces and torques a motile cell would encounter. A cell experiences a propulsion (***P***) attributed to the drive from its flagella, the weight of its own body which can be segregated into weight from its nucleus, lipids and cytoplasm, an upthrust due to its density being not equal to the surrounding fluid and a drag force attributed to the viscous effects. The relevant torques about its center of buoyancy will include one from nucleus and lipids (if they do not reside on the major axis of the cell), torque due to viscous drag and torque induced due to translational drag for asymmetric cells (*48*). With these considerations, we formulate the overall force and torque balance in each component as follows:

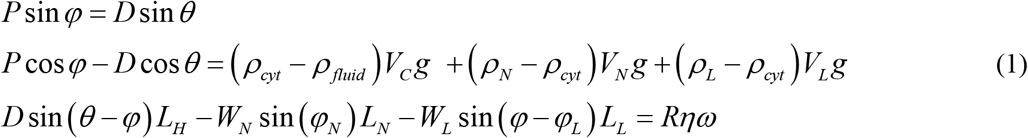

**Figure M1.**
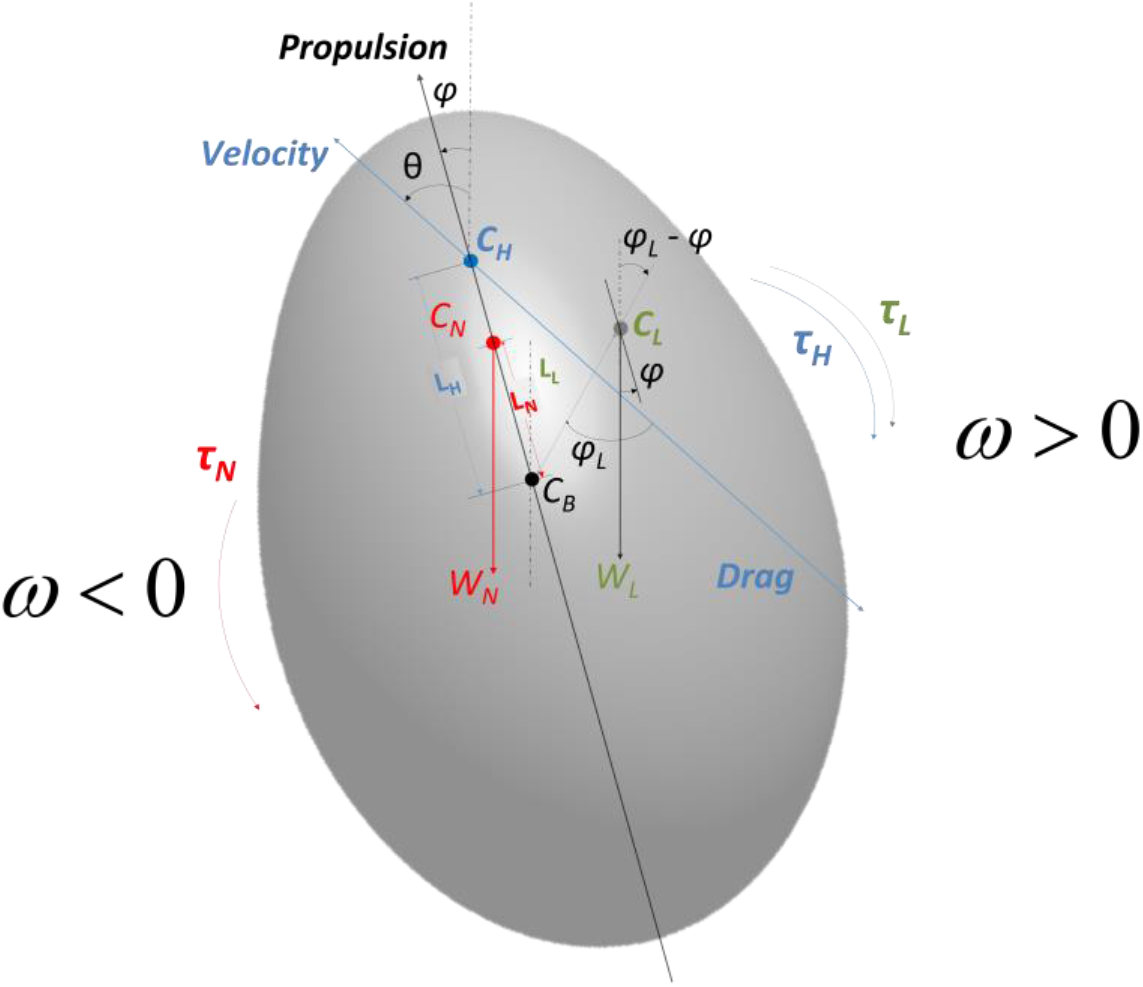
Schematics of the geometry of the single-cell of the phytoplankton along with the free body diagram of all forces and torques.

Here *ρ, L, V* and *η* denotes the density, distance from cell centroid, volume, and medium viscosity respectively. The subscripts *C*, *cyt, fluid, N, L* and *H* refers to the cell, cytoplasm, background medium, nucleus, lipids and the hydrodynamic center of the cell, respectively. *φ*_*N*_ is the angle between the direction of gravity (vertical line) and the line joining *C*_N_ and *C*_B_ while *φ*_*L*_ is the angle between the major axis and the line joining *C*_L_ and *C*_B_. *φ* is the angle of the cell propulsion axis (resultant flagellar motion) and the vertical axis, which is an unknown along with the propulsion force **P**. Such definitions of the angles make them independent of the initial angular position of the cell and dependent on *φ*, which comes as a part of our from of equation (1). Since we assume the center of gravity of the nucleus to lie on the major axis, an assumption experimentally very close to, therefore *φ*_*N*_ = *φ*. Realistically, a cell does not move exactly in the direction of propulsion and is assumed to move at an offset making an angle *θ*, a known parameter obtainable from experiments, with the vertical. In the above configuration, counter clockwise angular direction from the vertical line (direction of gravity) is assumed to be positive. The terms *W_L_* and *W_N_* are the weight of the lipids and the nucleus with respect to the cytoplasm. *D* is the drag force described below, *P* is the force of propulsion which is unknown and is obtained from solving equations set (1). The cell is described by the generic equation 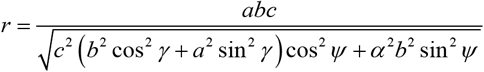 where *a, b* (*a>b*), *c (=b)*, *γ* ( 0 <*γ*< *2π*) and *ψ* (−*π*/2<*ψ*<*π*/2) respectively denotes the major axis length, semi-major axis length, minor axis length, azimuth angle and polar angle. The symmetric geometry implies that the hydrodynamic center of the cell is on the cell centroid (centre of buoyancy) and *L*_*H*_ vanishes (*48*). Finally, *R* represent the coefficient of hydrodynamic rotational resistance on the phytoplankton cell. To obtain the cell dimensions, we have fitted the phase contrast microscopy images to the cell profile using above equation and imposing *γ* = 0, thereby obtaining a 2D version of the parametric equation of the form 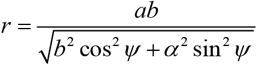 (*65*). The cell being assumed symmetric prolate ellipsoid, coefficient of resistance *R* and the drag, expressed as *D*_∥,⊥_ = 6*πηr*_*eq*_*UK*_∥,⊥_, where *U* is the translational velocity and *K* denotes the shape factor, are re-derived for our case (*66*, *67*) with *t=a/b*. The viscous torque drag can be estimated using *R* where *ω* is the angular velocity. All the meaning of the rest of the symbols is geometrically explained in the above model geometry. This set of three equations have three unknowns namely ***P, ϕ*** and ***ω***. The solution we are interested is in finding the angular rotation rate ***ω***. Thus, from all the values known from the experiments across different growth stage, we attempt to draw a phase plot (represented in Figure 4E-G) that reflects the value of the angular rotation rate as a function of the different possible position (varying *φ*_*L*_ and *L*_*L*_) a representative LD of dimensions corresponding to a particular growth stage can take within a cell. The phase plots highlight the stability of the cell that has contribution from various factors. With nucleus remaining very close to the geometric center, the dominant factor to impart the stability criterion to the strain S-1 cells is the lipid compartmentalization effects.

The active torque can now be estimated in a straight-forward manner from last equation of set (1) with unbalanced torque expression. Note that for any general ellipsoid, the drag force *D* applied by an arbitrary ellipsoid, with semi-axis lengths {k, m, n}, on the fluid when translating at speed *U* in the *k* direction, is 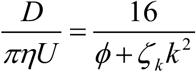, while the torque applied on a fluid due to rotation with rate *w* around the *k* direction is 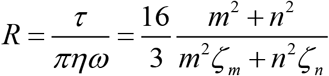 (*68*) where the variables in the relation is given by 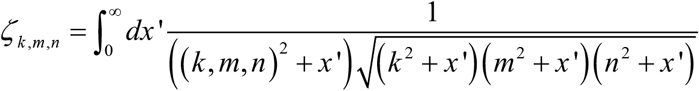 (with *k*, *m*, and *n* denoting individual relations for *ζ*_*k*_, *ζ*_*m*_ and *ζ*_*n*_, respectively) and 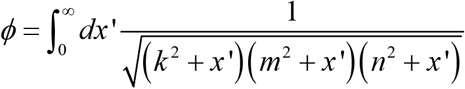. *R = R* for axially symmetric shape about the major axis. We have estimated the integrals using MATLAB and matched with the assymptotic analytical expression for axially symmetric spheroid (prolate). The active torque is plotted as a function of different ratios of the shortest axis (*c*) and major axis (*a*), different ratios of the lipid volume by cell volume and different ratios of the lipid distance from the cell geometric centre to major axis of the cell. All other factors for strain S-1 and S-2 are kept same for comparison of the required active torque.

### Reorientation analysis in the dynamic experimental setup

The swimming stability of *H*. *akashiwo* 3107 and 452 strains was quantified for two time points representative of exponential (72 h) and stationary (700 h) growth stages. Experiments were conducted in rectangular millifluidic chamber (10 mm x 3.3 mm x 2 mm) made of PPMA (Poly methyl methacrylate) with ~ 1.6 mm inlet and outlet circular holes located at the edges of chamber opposite to each other. The chamber was mounted via magnets on a circular plate which is, in turn, attached to a custom-made holding cage that include light diffuser (220 grits) and a red LED (630 nm wavelength). The different components in the cage are placed in a way to permit homogenous lit of the interested chamber area. One end of the entire cage is connected to the shaft of a stepper motor that permits the rotation of the whole setup as needed. The Nema 23 stepper motor was controlled automatically using a DM452T driver coupled to an Arduino uno board. The automation part comes from programming the Arduino to rotate by a specific degree for a specific duration. On the other end of the cage, a variable zoom lens system (zoom = 1.7x) was connected to a Grasshopper3 (Model: GS3-U3-41C6C-C) camera with a 1’’ sensor that permits to capture the entire vertical height of the chamber. The geometric center of the chamber, light source and diffuser were placed in coaxial position. For imaging, the focal plane was chosen visually to be in the middle of the experimental millifluidic chamber.

Prior to image acquisition, concentration of cells in the measured sample was adapted to permit single-cell tracking and quantification of corresponding reorientational stability (*48*). Due to low cell density, no dilution was required at 72 hours. During late stationary stages, the cell cultures were diluted in a glass vial supplemented with 3.4 mL of 0.45 μm filtered culture supernatant collected from the bottom of the experimental culture tube using a 120 mm needle and 2 mL syringe. 1.2 mL cell culture collected from the top 5 mm tube were gently added in the filtered supernatant volume to reach a 3.8x final dilution. The glass vial with the cell dilution was covered with parafilm with multiple holes done using small needle to allow airflow, and placed in the incubator at 22°C for 50 minutes to allow cells acclimatization. We conducted two biological replicates with three technical and three technical replicates, numbering a total of 18 replicates for each of the two strains analyzed. Millifluidic chamber was filled by gently pipetting suspension of phytoplankton cells (~180 μL) sampled from the top 0.5 cm culture followed by closing the inlet and outlet ports with a silicone plug and mounting it on the magnetic cage. A wait time of 20 minutes was given so that cells distributed in the most stable configuration in the millifluidic chamber. Afterward, a 180-degree rotation was applied to the chamber through the Arduino controlled automatic stepper motor (programmed in-house). Once cells swam to the middle of the chamber, we applied a series of three consecutive flips with intervals of 15, 12, 12 seconds for cells at 72 hours, and 25, 20, 20 seconds for cells at 700 hours. The different waiting periods were adapted in light of the growth stage-dependent reorientation speeds (speed at stationary stage is lower than speed at exponential stage, Figure 3). Videos were acquired at 16 frames per second. Phytoplankton cells were tracked using Image J plugin Mosaic Particle Tracker (2D/3D), and analyzed using in-house Python codes. We analyzed 160 frames and 220 frames for populations at 72 and 700 hours, respectively. The corresponding swimming times of 10 and 14 seconds were sufficient to capture multiple cell reorientation events in each replicate. Among the acquired trajectories, those which appear for less than 45 frames, or those which showed a net displacement less than 10 pixels (corresponding to 20 μm, one cell body length) were eliminated. The remaining trajectories were interpolated quadratically (smoothing). For single trajectory, angular velocity (ω) was obtained for each consecutive frame as a function of the instantaneous angular position (θ). Angular velocities were averaged for given θ value (ranging from −90 to 90 degrees, binned at intervals of 10 degrees). Any ω value greater than 0.5 rad/sec or less than −0.5 rad/sec was eliminated. Obtained ω values were fitted to a sinusoidal curve of the form A × sin(x), and the reorientation timescale was then obtained as 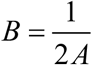.

**Figure M2.**
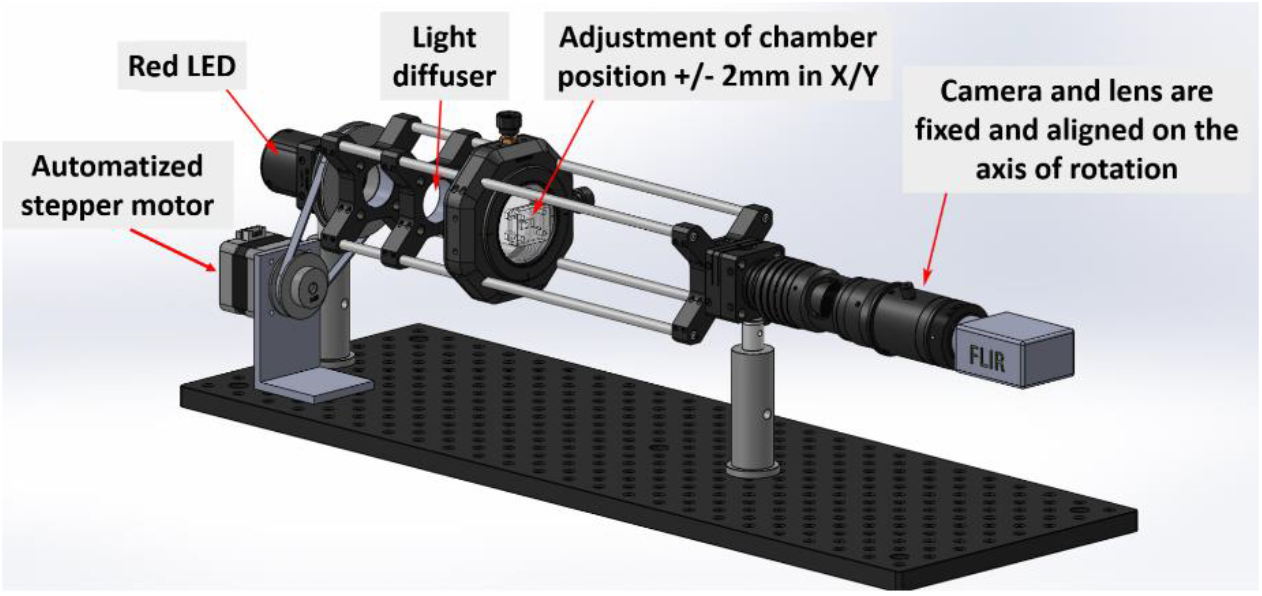
Schematic of the microscopic imaging system that is used to find the reorientation time scale of the phytoplankton population.

### Quantification of endogenous cellular stress

Endogenous reactive oxygen species (ROS) were quantified with the fluorescent stain CellROX Orange (Thermofisher, excitation/emission 545/565 nm) to determine impact of nutrient limitation on ROS production. CellROX Orange is a cell-permeable reagent non-fluorescent while in a reduced state and upon oxidation exhibits strong fluorogenic signal. *H*. *akashiwo* cells sampled at different nutrient regimes were incubated with 5 μM CellROX Orange for 30 minutes in the dark (400 μL cells with 0.6 μL CellROX ready-to-use stock solution). After incubation, cells were illuminated using green light ( ~ 545 nm) with an exposure time of 1/5, and an LED fluorescence intensity of 100%. The fluorescence readout was quantified over 21 seconds using fluorescence microscopy (Olympus CKX53 inverted microscope) supplemented with high-resolution color camera (Imaging Source, DFK33UX265). For single cell, the fluorescence intensity is increased over time; the magnitude of fluorescence intensity before cell lysis corresponded to the maximum ROS accumulation in the cell. For quantification, acquired single-cell fluorescence image was analyzed with ImageJ Z-stack layer to extract time-dependent signal intensity variation. Stress accumulation rate, quantified as fluorescence intensity signal, was integrated over the first 10 seconds of acquisition.

## Notes

**Funding:** This work was supported by the ATTRACT Investigator Grant (Grant no. A17/MS/11572821/MBRACE), the PRIDE Doctoral Training Unit (project PRIDE19/14063202/ACTIVE), and the AFR Grant (Grant no. 13563560) of the Luxembourg National Research Fund. The work received generous support from the Human Frontier Science Program Cross Disciplinary Fellowship (to J.D., LT000368/2019-C), and the Swiss National Science Foundation Early Postdoc Mobility (to F.D., project number: P2GEP3_184481).

### Competing Interest Statement

The authors have declared no competing interest.

